# A PRDM16-CtBP1/2 Complex Interacts with HDAC1/2 to Regulate Transcriptional Programs of Neurogenesis and Guide Cortical Neuron Migration

**DOI:** 10.1101/2024.11.17.624022

**Authors:** Sophie Warren, Bader El Farran, Sungyun Kang, Adhyeta Choudhuri, Sen Xiong, Volker P. Brendel, José-Manuel Baizabal

**Affiliations:** Department of Biology, Indiana University, Bloomington, IN, 47405, USA; Department of Computer Science, Indiana University, Bloomington, IN, 47405, USA

## Abstract

Chromatin regulation of transcriptional enhancers plays a central role in cell fate specification and differentiation. However, how the coordinated activity of transcription factors and chromatin-modifying enzymes regulates enhancers in neural stem cells (NSCs) and dictates subsequent stages of neuronal differentiation and migration is not well understood. The histone methyltransferase PRDM16 is expressed in NSCs of the developing mouse and human cerebral cortex and is essential for determining the position of upper-layer cortical neurons. Here, we report that PRDM16 interacts with C-terminal binding protein 1 (CtBP1) and CtBP2 to control the transcriptional programs of cortical neurogenesis and regulate upper-layer neuron migration. PRDM16 and CtBP1/2 co-regulate enhancers by interacting with histone deacetylase 1 (HDAC1) and HDAC2, and lysine-specific demethylase 1 (LSD1). In addition, our results suggest that the CCCTC-binding factor CTCF plays a key role in recruiting CtBP1/2 to cortical enhancers. These findings underscore that reduced interactions between PRDM16 and ubiquitous chromatin regulators may contribute to neurodevelopmental deficits in patients with *PRDM16* haploinsufficiency.

## INTRODUCTION

The remarkable complexity of the mammalian cerebral cortex originates during embryonic development from a population of neural stem cells (NSCs) also known as radial glia (Casingal et al., 2022). Cortical NSCs reside in the ventricular zone (VZ), where they differentiate into intermediate progenitors that migrate into the subventricular zone (SVZ) and give rise to deep-layer neurons at early stages and upper-layer neurons at later stages of embryonic neurogenesis (Kwan et al., 2012)(Di Bella et al., 2024). Newborn neurons migrate from the VZ/SVZ to the cortical plate guided by the long basal processes of NSCs that span across the cortex (Noctor et al., 2001)(De Juan Romero and Borrell, 2015). The precise control of neuronal migration is critical to establishing cortical organization into six neuronal layers, which produce the circuits that control behavior and higher cognition in humans.

Previous studies have elucidated a critical role for multiple transcription factors and chromatin-modifying enzymes in regulating cortical neurogenesis, neuronal fate specification, and migration (Kwan et al., 2012)(Amberg et al., 2019). A subset of chromatin-modifying enzymes influences gene expression by “writing” or “erasing” covalent histone modifications, such as methylation and acetylation (Jambhekar et al., 2019). Histone methylation and acetylation are dynamically regulated at enhancers during cortical neurogenesis and play a pivotal role in controlling cell-type-specific gene expression (Amberg et al., 2019)(Albert et al., 2017). The emerging concept of epigenetic pre-patterning postulates that histone modifications promote chromatin looping interactions between distal enhancers and their target genes, resulting in topology changes that are permissive to gene activation upon differentiation (Albert and Huttner, 2018). However, how the interplay between transcription factors and chromatin-modifying enzymes establishes chromatin modifications at enhancers in NSCs, enabling subsequent stages of neuronal differentiation, migration, and maturation, remains an open question.

PRDM16 (PR domain-containing 16) is a histone methyltransferase with multiple regulatory roles in neural cells, hematopoietic cells, myocytes, and adipocytes (Chi and Cohen, 2016). In cultured fibroblasts, PRDM16 catalyzes the mono-methylation of lysine 9 on histone H3 (H3K9me1), leading to transcriptional repression and chromatin compaction (Pinheiro et al., 2012). In contrast, PRDM16 promotes histone H3 lysine 4 mono- and di-methylation (H3K4me1 and H3K4me2), resulting in transcriptional activation in hematopoietic cells (Zhou et al., 2016). PRDM16 has two zinc finger domains and a repressor domain that mediates the interaction with other transcriptional regulators (Chi and Cohen, 2016)(Hohenauer and Moore, 2012). In the developing mouse and human cerebral cortex, *PRDM16* is specifically expressed in NSCs and regulates progenitor proliferation, neurogenesis, and the migration of projection neurons into upper cortical layers (Inoue et al., 2016)(Baizabal et al., 2018)(Garcıá et al., 2020)(He et al., 2021)(Suresh et al., 2024). PRDM16 activity in cortical NSCs controls histone acetylation and chromatin accessibility at developmental enhancers, thereby promoting the activation of genes associated with neurogenesis and repressing the premature expression of neuronal differentiation and migration genes (Baizabal et al., 2018)(He et al., 2021). Notably, *PRDM16* haploinsufficiency in humans is associated with 1p36 deletion syndrome and may result in reduced cortical folding, microcephaly, seizures, and intellectual disability—highlighting the importance of *PRDM16* in human brain development (Jordan et al., 2015)(Jacquin et al., 2023)(Suresh et al., 2024).

In this study, we sought to identify PRDM16 transcriptional partners and determine their functional cooperation in regulating corticogenesis. Our results indicate that PRDM16 forms a complex with the transcriptional regulator C-terminal binding protein 1 (CtBP1) and CtBP2 (collectively referred to as CtBP1/2). PRDM16 and CtBP1/2 bind common enhancer regions in the embryonic cortex and cultured NSCs, and co-regulate transcriptional programs of neurogenesis and neuronal migration. Accordingly, silencing of *Ctbp1/2* in the embryonic cortex results in impaired migration and ectopic upper-layer neurons in the postnatal brain. PRDM16 and CtBP1/2 interact with histone deacetylase 1 (HDAC1) and HDAC2 in cultured NSCs. In the embryonic cortex, PRDM16 and CtBP2 co-regulate enhancers with HDAC1 and lysine demethylase 1 (LSD1). Our data suggests that CtBP1/2 transcriptional complexes are recruited to genomic loci through an interaction with CTCF (CCCTC-binding factor). Together, our findings provide insights into how transcriptional and chromatin regulators collaborate to control enhancers, thereby facilitating cortical neurogenesis, and subsequent steps of neuronal differentiation.

## RESULTS

### CtBP1/2 interact with PRDM16 and control cortical neuron migration

To identify PRDM16-interacting partners, we generated primary cultures of NSCs from the embryonic day 14.5 (E14.5) cerebral cortex. These cultures consist of a relatively homogeneous monolayer of proliferating Ki67^+^ cells **(Fig. S1A)**. The majority of cortical cells expanded *in vitro* are PRDM16^+^ and NESTIN^+^, confirming their NSC identity **(Fig. 1A)** (Noctor et al., 2001)(Baizabal et al., 2018). Immunoprecipitation (IP) of PRDM16 in cultured NSCs followed by mass spectrometry and western blot (WB) identified a strong interaction with CtBP1 and CtBP2 **(Fig. 1B, S1B)**. Conversely, CtBP1/2 IP confirmed an interaction with PRDM16 **(Fig. S1C)**. CtBP1 and CtBP2 are primarily co-repressors that form transcriptional complexes with multiple chromatin-modifying enzymes (Shi et al., 2003)(Stankiewicz et al., 2014). *CtBP1* and *CtBP2* are broadly expressed in the embryonic cortex, including the VZ **(Fig. 1C)**, where *Prdm16* is expressed (Inoue et al., 2016)(Baizabal et al., 2018). Hence, the interaction between PRDM16 and CtBP1/2 is specific to NSCs during corticogenesis.

**Fig. 1.**
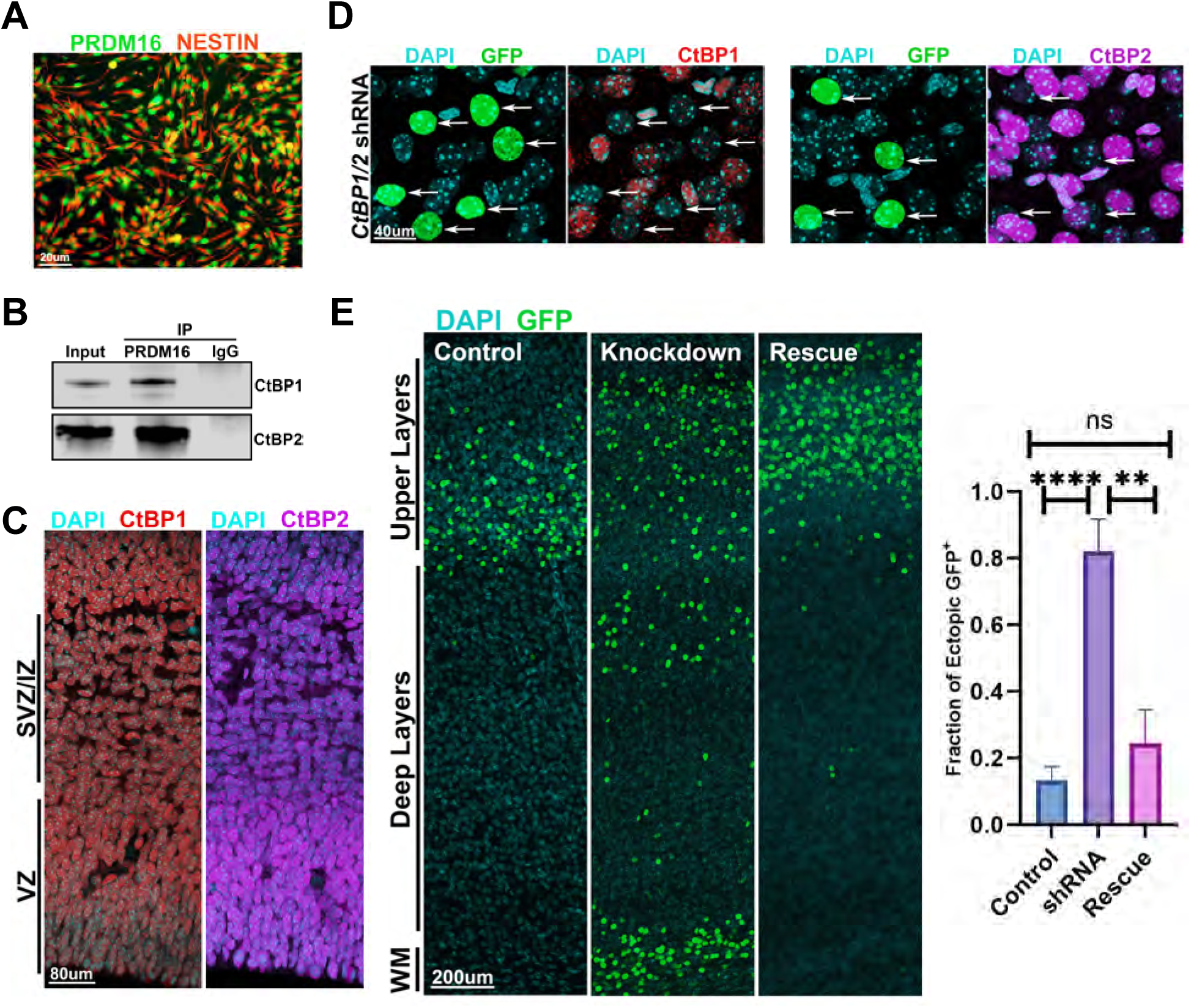
CtBP1/2 Interact with PRDM16 and control neuronal migration. (A) Primary culture of PRDM16^+^/NESTIN^+^ NSCs isolated from the E14.5 cortex. (B) Immunoprecipitation (IP) of PRDM16 followed by western blot indicates interaction with CtBP1 and CtBP2 in NSCs. An unspecific IgG was used as a negative control. (C) Immunostaining shows detection of CtBP1/2 in NSCs at the ventricular zone (VZ), as well as in differentiating cells at the subventricular zone (SVZ) and intermediate zone (IZ). (D) Knockdown of *Ctbp*1/2 by *in-utero* electroporation of shRNAs results in strong depletion of CtBP1/2 in GFP^+^ electroporated cells (arrows). (E) *In-utero* electroporation of *Ctbp1/2* shRNAs at E14.5 followed by the analysis of neuronal migration at P5 indicates a significant increase of GFP^+^ cells in deep layers and adjacent to the white matter (WM) in comparison to controls. In the rescue condition, *Ctbp1/2* shRNAs were co-electroporated with plasmids carrying the shRNA-resistant coding sequences of *Ctbp1/2*, resulting in a significant decrease of ectopic GFP^+^ cells in deep layers. Data represent mean ± SD, statistical analysis is unpaired Student’s t-test (**p < 0.01, ****p < 0.001, ns: not significant).

Previous studies have shown that CtBP2 regulates cortical neurogenesis, whereas *CtBP2* overexpression impairs cortical neuron migration (Karaca et al., 2020)(Wang et al., 2019). To test the function of *CtBP1* and *CtBP2* in the developing cortex, we simultaneously knocked down both genes by *in-utero* electroporation (IUE) of short-hairpin RNAs (shRNAs) during upper-layer neurogenesis at E14.5. To follow the fate of electroporated neurons, we co-injected a plasmid encoding Green Fluorescent Protein (GFP). We observed a strong depletion of CtBP1 and CtBP2 in most GFP^+^ neurons **(Fig. 1D)**. Analysis of cortical neuron position at postnatal day 5 (P5) revealed that *CtBP1/2* silencing results in many electroporated neurons ectopically located in deep-layers and near the white matter tracts in comparison to controls **(Fig. 1E)**. The ectopic neurons displayed *Brn2* expression, confirming their upper-layer neuron identity **(Fig. S1D)**. To test the specificity of the observed migration defects, we performed a rescue experiment by co-electroporating shRNAs targeting *CtBP1/2* together with shRNA-resistant coding sequences of *CtBP1/2*. We confirmed strong expression of *CtBP1/2* in GFP^+^ cells in the rescue condition **(Fig. S1E)**. The rescue of *Ctbp1/2* expression in shRNA-electroporated neurons led to a significant reduction in the number of ectopic neurons **(Fig. 1E)**. The impact of *CtBP1/2* silencing on upper-layer neuron migration is strikingly similar to the one observed upon silencing *Prdm16* in the embryonic cortex (Baizabal et al., 2018), suggesting that the interaction between PRDM16 and CtBP1/2 in NSCs influences cortical neuron migration.

### CtBP1/2 regulate transcriptional programs of neurogenesis and neuronal migration

Previous work indicates that *CtBP1/2* are ubiquitously expressed throughout the developing mammalian embryo (Hildebrand and Soriano, 2002). To test whether CtBP1/2 regulate broad or neural-specific gene expression programs, we infected cortical NSCs *in vitro* with lentiviruses carrying shRNAs against *CtBP1* and *CtBP2*, resulting in simultaneous depletion of CtBP1 and CtBP2 **(Fig. 2A)**. We then performed RNA-seq to identify differentially expressed genes between *CtBP1/2* knocked down NSCs and control cultures infected with a scrambled shRNA. We initially confirmed that *Ctbp1* and *Ctbp2* were robustly silenced in all four RNA-seq replicates in comparison to controls **(Fig. S2A)**. Despite the reported predominant role of CtBP1/2 as transcriptional co-repressors in non-neural tissues (Stankiewicz et al., 2014), we observed a similar number of both upregulated and downregulated genes in *CtBP1/2* knocked down NSCs **(Fig. 2B)** (Supplementary file 1). Gene ontology analysis indicated that CtBP1/2 activity in cortical NSCs primarily regulates genes associated with neural development, including cell migration and neurogenesis **(Fig. 2C)**.

**Fig. 2.**
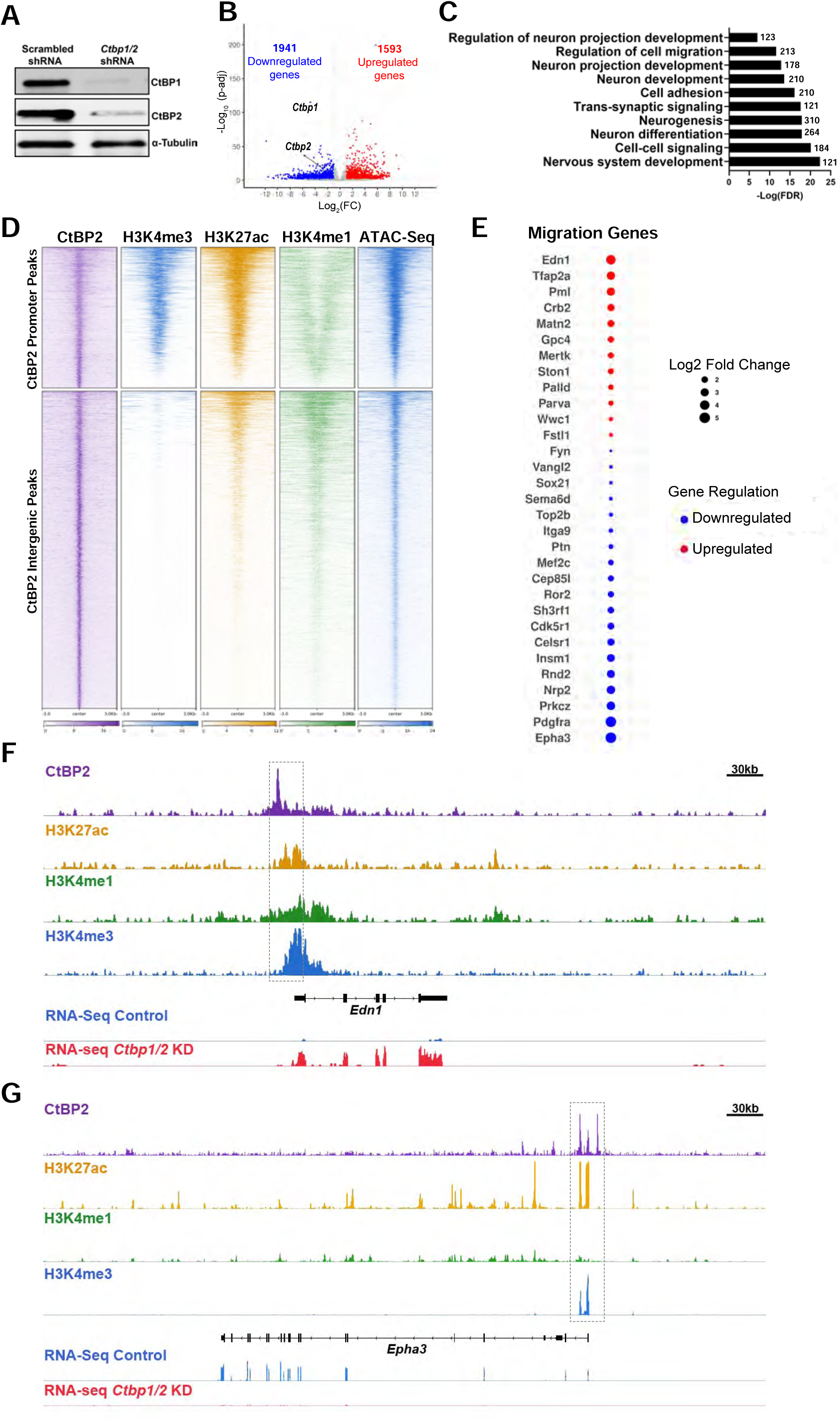
CtBP1/2 control transcriptional programs of cortical neurogenesis and migration. (A) Cortical NSCs co-infected with lentiviruses carrying *Ctbp1/2* shRNAs exhibited strong depletion of CtBP1 and CtBP2 by western blot. The loading control is α-Tubulin. (B) Volcano plot showing differentially expressed genes between NSCs infected with *Ctbp1/2* shRNAs and scrambled controls. *Ctbp1* and *Ctbp2* downregulation are indicated in the plot. RNA-seq data were generated from four biological replicates. (C) Gene ontology analysis of differentially expressed genes between the *Ctbp1/2* knocked down and control NSCs. (D) Heatmap of CtBP2 binding sites at promoters and intergenic regions. The overlaps with active promoters (H3K4me3), active/poised enhancers (H3K27ac/H3K4me1), and highly accessible chromatin regions (ATAC-seq) in the embryonic cortex are also shown. (E) Dot-plot of neuronal migration genes directly regulated by CtBP2 that display differential expression in the *Ctbp1/2* knocked down NSCs. (F) Genome tracks showing CtBP2 binding to the promoter of the cell migration gene *Edn1*. The RNA-seq tracks indicate *Edn1* upregulation in the *Ctbp1/2* knockdown (KD), suggesting that CtBP2 directly repress this gene. (G) Genome tracks displaying CtBP2 binding to the promoter of the migration gene *Epha3*. The RNA-seq tracks suggest that *Epha3* is directly activated by CtBP2.

To investigate the regulatory mechanisms by which CtBP1/2 control neural gene expression, we performed chromatin IP with high-throughput sequencing (ChIP-seq) for CtBP2 in the E14.5 cortex and cultured NSCs. By intersecting the *in vivo* and *in vitro* ChIP-seq datasets, we identified 12020 reproducible CtBP2 binding sites in NSCs which are also validated in the embryonic cortex **(Fig. S2B)** (Supplementary file 2). A subset of 3012 CtBP2-binding regions are located in active promoters of the E14.5 cortex, as indicated by the overlap with H3K4me3 **(Fig. 2D)**. A distinct subset of 2093 CtBP2 binding sites overlapped with H3K27ac and H3K4me1, suggesting an association with active enhancers **(Fig. 2D)**. Furthermore, CtBP2 binding sites displayed high chromatin accessibility, as determined by the Assay for Transposase-Accessible Chromatin with sequencing (ATAC-seq) **(Fig. 2D)** (Gorkin et al., 2020). These results indicate that CtBP2 binds dynamically regulated promoters and enhancers in the embryonic cortex.

Next, we sought to identify genes directly regulated by CtBP2-bound enhancers in NSCs of the embryonic cortex. To achieve this goal, we used previously reported high-throughput chromosome conformation capture (Hi-C) datasets from the developing mouse cortex (Bonev et al., 2017). These datasets identified chromatin looping interactions between enhancers and promoters in cortical progenitors and neurons (Bonev et al., 2017). Each Hi-C enhancer-promoter pair (EPP) is associated with one gene, while genes regulated by multiple enhancers and linked to several EPPs (Bonev et al., 2017). We intersected CtBP2 binding sites with annotated promoters and Hi-C enhancers to generate a compiled list of proximal and distal gene targets. Altogether, 2368 genes are directly regulated by CtBP2 in the embryonic cortex, out of which 1654 genes are regulated through promoter binding and 807 genes by enhancer binding (Supplementary file 2). To identify gene targets for which CtBP1/2 binding is critical, we intersected the list of predicted regulated genes with dysregulated genes in the *Ctbp1/2* knocked down NSCs. We found that CtBP2 is necessary to directly control the expression of a group of genes linked to neurogenesis and cell migration **(Fig. 2E, S2C)**. Surprisingly, CtBP2 is critical for directly activating 239 genes and silencing 127 genes, suggesting that CtBP2 functions as both an activator and repressor in the developing cortex **(Fig. 2F, G)** (Supplementary file 2). Collectively, our results suggest that CtBP2 binding at promoters and enhancers activates and represses genes associated with neurogenesis and neuronal migration during corticogenesis.

### PRDM16 and CtBP2 bind to an overlapping subset of enhancers to regulate gene expression

PRDM16 and CtBP1/2 interact in cortical NSCs and both dictate neurogenesis and neuronal migration **(Fig. 1B, E)** (Baizabal et al., 2018)(Karaca et al., 2020). These data suggest that PRDM16 and CtBP2 collaborate to regulate gene transcription in NSCs. To test this possibility, we initially sought to identify the genes directly regulated by PRDM16 in the embryonic cortex. To perform this analysis, we combined annotated promoters and Hi-C EPPs with PRDM16 binding sites, resulting in the identification of 2213 putative gene targets **(Fig. S3A)** (Supplementary file 3). A subset of 185 predicted targets were dysregulated in NSCs and intermediate progenitors of the *Prdm16* knockout (KO) embryonic cortex (Supplementary file 3) (Baizabal et al., 2018). Our analysis indicates that PRDM16-bound enhancers and promoters directly regulate genes associated with cell migration and neurogenesis **(Fig. S3B)**.

We assessed the overlap between PRDM16 and CtBP2 binding sequences, which resulted in 1023 sites bound by both factors in cortical NSCs **(Fig. 3A)**. The majority of PRDM16 and CtBP2 overlapping binding sites are located within enhancers **(Fig. S3C)**. We intersected the predicted gene targets (based on promoter binding and Hi-C EPP overlaps) of PRDM16 and CtBP2 (2213 versus 2368 genes, respectively) and found 793 genes directly co-regulated by both factors **(Fig. 3B)** (Supplementary file 4). PRDM16 and CtBP2 bind to a group of Hi-C-identified enhancers that are predicted to regulate most of the 793 gene targets **(Fig. 3C)** (Bonev et al., 2017). Furthermore, we found that 169 out of 793 co-regulated genes require PRDM16, CtBP2, or both for normal expression in cortical cells (Supplementary file 4). Gene ontology analysis of PRDM16 and CtBP2 co-regulated genes indicate enrichment in neurogenesis and cell migration **(Fig. 3D)**. A subset of predicted gene targets associated with neurogenesis and cell migration are distally regulated by PRDM16- and CtBP2-bound enhancers **(Fig. 3E, F)**. Together, these results suggest that PRDM16 and CtBP1/2 cooperate in NSCs to control gene transcription, primarily through enhancer regulation.

**Fig. 3.**
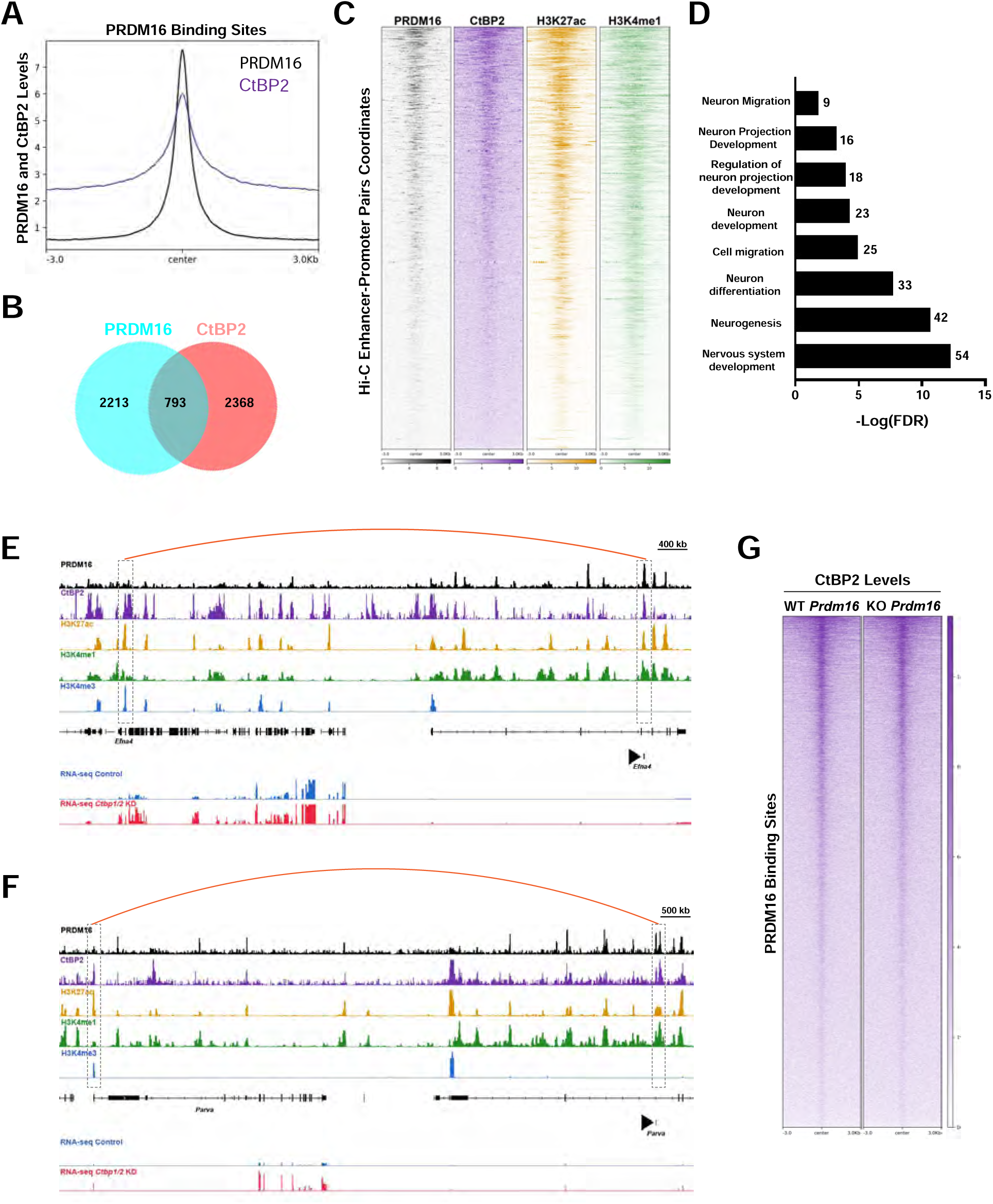
PRDM16 and CtBP2 co-regulate enhancers and gene targets in the embryonic cortex. (A) Projection profile showing PRDM16 and CtBP2 binding levels within PRDM16-regulated sequences in NSCs of the E14.5 cortex. (B) Venn diagram showing the number of genes directly regulated by PRDM16, CtBP2, or both in the embryonic cortex. (C) Heatmaps showing PRDM16 and CtBP2 binding within enhancers with identified gene targets. The heatmap coordinates represent the enhancer regions of enhancer-promoter pairs previously identified by Hi-C in the embryonic cortex (Bonev et al., 2017). (D) Gene ontology analysis for genes co-regulated by PRDM16 and CtBP2. (E, F) The genome browser tracks show two examples of enhancers bound by PRDM16 and CtBP2 (right dashed-line boxes) that are predicted to interact with the promoters (left dashed-line boxes) of the neurogenesis gene *Efna4* (E) and the migration gene *Parva* (F). The RNA-seq tracks show *Efna4* and *Parva* upregulation in the *Ctbp1/2* knockdown (KD). Arrowheads indicate the BED file coordinates of the intersection between PRDM16, CtBP2, H3K27ac, and Hi-C enhancer-promoter pairs. The predicted gene target of each enhancer is indicated below the arrowhead. The curve lines indicate the predicted long-range interactions between enhancers and promoters. (G) Heatmaps displaying CtBP2 binding in *Prdm16* WT and *Prdm16* KO NSCs *in vitro*. The heatmap coordinates represent PRDM16-regulated sequences.

Previous studies in adipocytes suggest that PRDM16 recruits CtBP1/2 to white fat selective genes, leading to transcriptional repression (Kajimura et al., 2008). To test whether PRDM16 recruits CtBP2 to neural enhancers, we generated *Prdm16* wild-type (WT) and KO cultures of NSCs from the E14.5 cortex **(Fig. S3D)**. We then performed quantitative ChIP-seq for CtBP2 in *Prdm16* WT and KO cultures. The normalized binding enrichment of CtBP2 within sequences co-regulated with PRDM16 in WT NSCs did not decrease in *Prdm16* KO NSCs **(Fig. 3G)**. Of note, the absence of PRDM16 leads to dysregulation of this subset of enhancers in the embryonic cortex (Baizabal et al., 2018). Therefore, these results suggest that CtBP1/2 remain associated with enhancers in the absence of PRDM16 but fail to properly regulate them, leading to impaired gene expression, and neuronal migration defects.

### PRDM16 and CtBP1/2 interact with HDAC1/2 and co-regulate enhancers in cortical cells

In the embryonic cortex, PRDM16 primarily promotes enhancer silencing, partly through promoting H3K27 deacetylation (Baizabal et al., 2018). However, the mechanisms by which PRDM16 controls histone acetylation at cortical enhancers remain unknown. Previous reports indicate that CtBP1/2 interact with HDAC1/2 to promote transcriptional silencing (Shi et al., 2003). Hence, we tested whether PRDM16 co-interacts with CtBP1/2 and HDAC1/2 in cortical cells. Using PRDM16 IP followed by WB, we detected HDAC1 and HDAC2 as interacting partners of PRDM16 in cultured NSCs **(Fig. 4A)**. Accordingly, 1168 PRDM16 binding sites overlap with HDAC1 in the E14.5 cortex **(Fig. 4B)**. Moreover, we found that CtBP1/2 also interact with HDAC1 and HDAC2 in cortical NSCs *in vitro* **(Fig. S4A)**. To examine whether PRDM16 binds HDAC1/2 through CtBP1/2, we introduced two point mutations in the PLDLS motif of PRDM16 to inhibit its interaction with CtBP1/2 (Kajimura et al., 2008). Consistent with a previous report, these point mutations completely abolished the interaction between PRDM16 and CtBP1/2 *in vitro* **(Fig. S4B)** (Kajimura et al., 2008). We then used lentiviruses to introduce the WT and mutant *Prdm16* coding sequences into primary cultures of *Prdm16* KO NSCs generated from E14.5 cortices. The interaction between PRDM16 and HDAC1/2 in NSCs was lost in the absence of PRDM16 binding to CtBP1/2 **(Fig. S4C)**. Our data indicate that PRDM16 interacts with HDAC1/2 through CtBP1/2 and these factors cooperate to regulate enhancers in NSCs of the embryonic cortex.

**Fig. 4.**
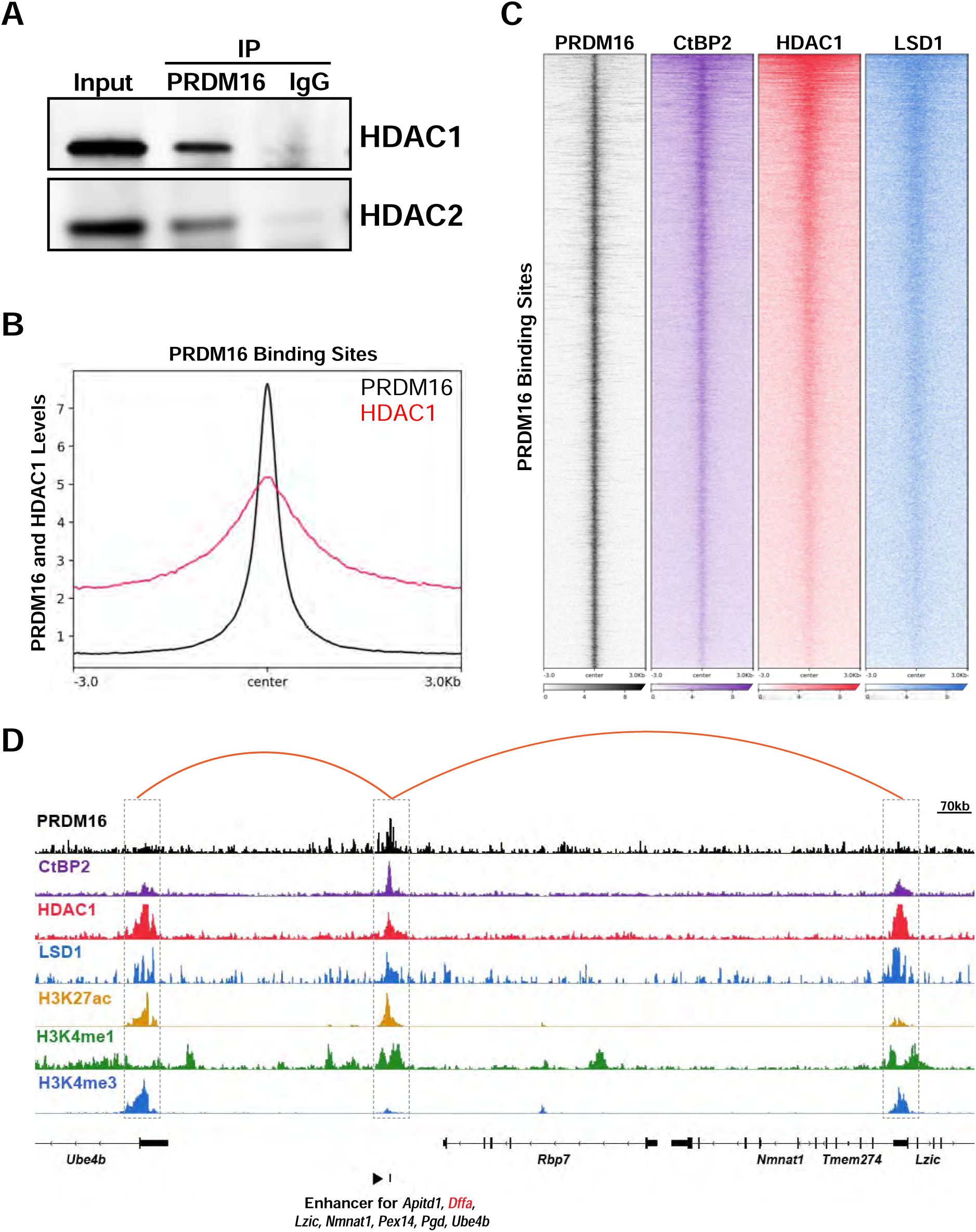
PRDM16 and CtBP2 co-regulate enhancers with HDAC1 and LSD1 in the embryonic cortex. (A) Immunoprecipitation (IP) of PRDM16 shows interaction with HDAC1 and HDAC2 in cortical NSCs *in vitro*. An unspecific IgG was used as a negative control. (B) Profile projection of PRDM16 and HDAC1 levels within PRDM16-regulated sequences. (C) Heatmaps showing PRDM16, CtBP2, HDAC1, and LSD1 levels within PRDM16-bound sequences in the embryonic cortex. (D) The genome tracks show an enhancer bound by PRDM16, CtBP2, HDAC1, and LSD1 (middle dashed-line box) that is predicted to interact with the promoter of seven genes, three of which are shown in the tracks (*Ube4b*, *Lzic*, and *Nmnat1*). The arrowhead indicates the BED file coordinates of the intersection between PRDM16, CtBP2, HDAC1, LSD1, and enhancer-promoter pairs previously identified by Hi-C in the embryonic cortex (Bonev et al., 2017). The seven genes regulated by the enhancer are indicated below the arrowhead. *Dffa* (red letters) is dysregulated in the *Ctbp1/2* knocked down NSCs (not shown). The long-range interactions between the enhancer and the promoters of *Ube4b*, *Lzic*, and *Nmnat1* (lateral dashed-line boxes) are indicated with curve lines. The histone marks show the relative enrichment of H3K4me3/H3K27ac at promoters and H3K4me1/H3K27ac at enhancers.

We next explored a potential co-regulatory role between PRDM16 and LSD1, a chromatin-modifying enzyme that interacts with CtBP1/2 and mediates enhancer silencing through H3K4me1 demethylation (Whyte et al., 2012)(Ray et al., 2014)(Zeng et al., 2016). We found that PRDM16 binds to a common subset of enhancers with CtBP2, HDAC1, and LSD1 **(Fig. 4C)**. Additionally, we identified a subset of distal gene targets regulated by enhancers bound by PRDM16, CtBP2, HDAC1, and LSD1 **(Fig. 4D)**. Together, our results suggest that interaction between PRDM16 and broadly expressed transcriptional regulators controls enhancer activity in cortical NSCs, a process critical for determining the positioning of cortical neurons.

### CTCF is a candidate factor for recruiting CtBP1/2 to cortical enhancers

In non-neural tissues and cell lines, CtBP1/2 are recruited to specific regulatory sequences by DNA-bound transcription factors (Chi and Cohen, 2016)(Stankiewicz et al., 2014). To investigate which transcription factors are potentially involved in recruiting CtBP1/2 to DNA in cortical NSCs, we performed DNA motif analysis using CtBP2 binding sites and PRDM16-CtBP2 binding sites. We found that the top consensus DNA motifs are bound by CTCF and BORIS (also known as CCCTC-Binding Factor Like or CTCFL) in both groups **(Fig. 5A)**. Motif analysis within PRDM16 binding sites alone does not result in a strong enrichment of CTCF motifs (Baizabal et al., 2018)(Garcıá et al., 2020). Hence, CTCF is a strong candidate for recruiting CtBP1/2 complexes to regulatory sequences. The gene encoding CTCF displays broad expression, whereas BORIS is restricted to germ cells (Lobanenkov and Zentner, 2018). CTCF mediates long-range enhancer-promoter interactions by promoting chromatin looping (Dehingia et al., 2022)(Sun et al., 2022). Consistent with our motif analysis, CtBP2 exhibits a substantial overlap with CTCF binding sites in the E14.5 brain **(Fig. 5B)** (Stamatoyannopoulos et al., 2012). Moreover, PRDM16 binding overlaps with CTCF mainly within sequences showing high levels of CtBP2 binding **(Fig. 5C)**. PRDM16, CtBP2, and CTCF binding at a subset of enhancers regulate distal gene targets through long-range interactions **(Fig. 5D, E)** (Bonev et al., 2017). These results suggest that CTCF recruits CtBP1/2 complexes at regulatory sequences, thereby controlling the transcriptional programs of cortical neurogenesis and neuronal migration.

**Fig. 5.**
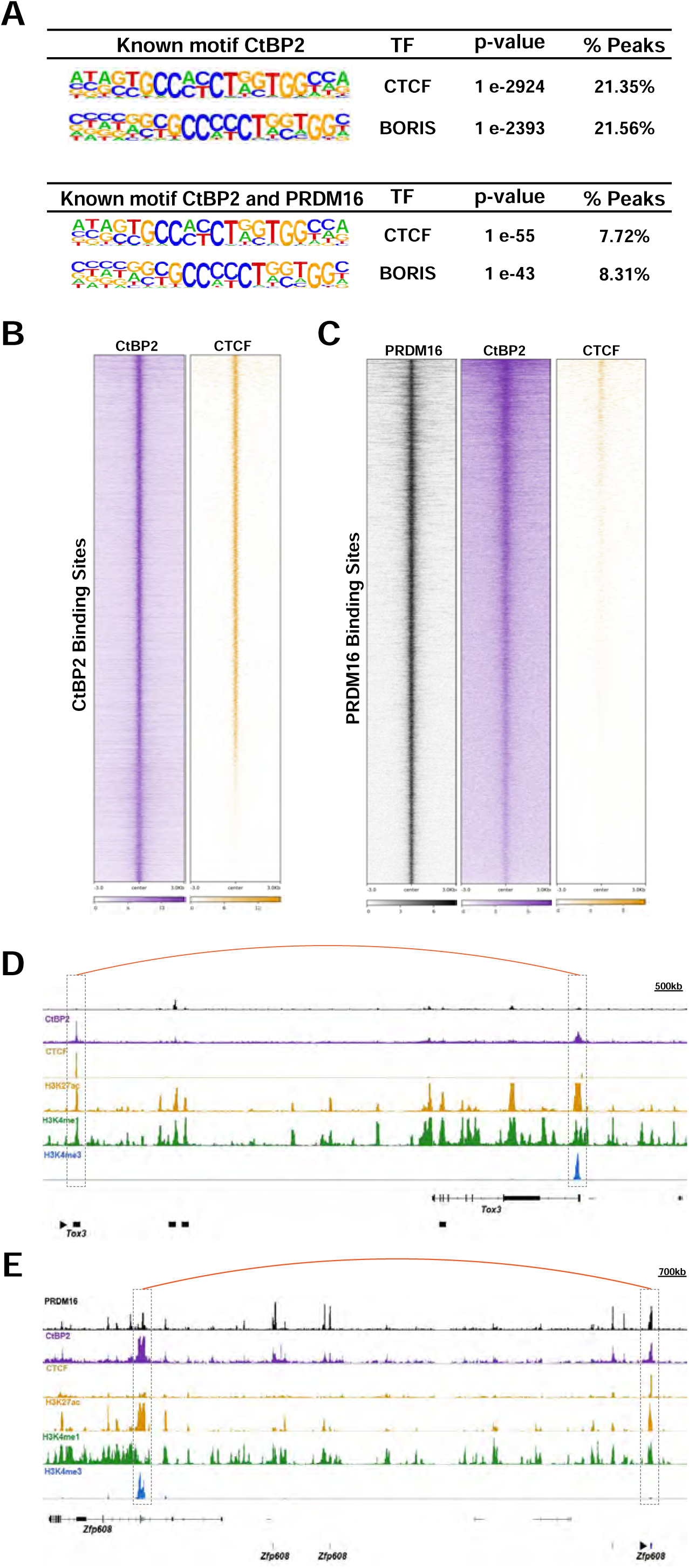
CTCF is a candidate recruitment factor for CtBP2 in the embryonic cortex. (A) Motif analysis within CtBP2 binding sites reveals significant enrichment of CTCF and BORIS (CTCFL) motifs. Accordingly, the sequences co-bound by PRDM16 and CtBP2 in cortical NSCs also show an enrichment of CTCF and BORIS motifs. (B) Heatmap showing a substantial overlap between CtBP2 and CTCF binding sites in the embryonic brain. (C) Heatmap showing that sequences with high levels of PRDM16 and CtBP2 binding also display a strong enrichment of CTCF binding (top of the heatmap). (D) The genome tracks display an enhancer bound by CtBP2 and CTCF (left dashed-line box) that interacts with the *Tox3* promoter (right dashed-line box). (E) The genome tracks show an enhancer co-regulated by PRDM16, CtBP2, and CTCF (right dashed-line box), which regulates the promoter of *Zfp608* (left dashed-line box). The curve lines in panels D and E indicate the long-range enhancer-promoter interactions predicted by Hi-C (Bonev et al., 2017). The arrowheads indicate the BED file coordinates of the intersection between CtBP2, CTCF, PRDM16, and Hi-C enhancers. The names of the distally regulated genes are indicated below the arrowheads. The histone marks show the relative enrichment of H3K4me3/H3K27ac at promoters and H3K4me1/H3K27ac at enhancers.

## DISCUSSION

In this study, we discovered an interaction between PRDM16 and CtBP1/2 in cortical NSCs and a critical role for CtBP1/2 in neuronal migration. Previous work has shown that *Ctbp2* knockdown and overexpression impair cortical neuron migration (Karaca et al., 2020)(Wang et al., 2019). However, the effects of knocking down *Ctbp2* on cortical neuron migration were analyzed during embryonic stages, leaving the possibility that the observed defects reflect a migration delay or an indirect effect due to a cell fate switch (Karaca et al., 2020). Conversely, overexpression of *Ctbp2* above physiological levels might not reflect the normal function of the endogenous CtBP2 at basal levels (Wang et al., 2019). Moreover, CtBP1 and CtBP2 are highly homologous proteins and likely play redundant roles (Hildebrand and Soriano, 2002). Hence, we decided to silence *Ctbp1* and *Ctbp2* simultaneously and perform the analysis of neuronal positioning at P5, when cortical neurons have completed their migration (Kwan et al., 2012). The silencing of *Ctbp1/2* led to impaired upper-layer neuron migration, which is strikingly similar to the migration defect observed in *Prdm16* KO embryonic cortex (Baizabal et al., 2018)(He et al., 2021). Hence, our results suggest that the interaction between PRDM16 and CtBP1/2 influences neuronal migration.

Previous work has shown that CtBP1/2 regulate progenitor proliferation and neurogenesis in the embryonic cortex and spinal cord (Karaca et al., 2020)(Dias et al., 2014). However, the mechanisms by which CtBP1/2 control gene expression in neural cells are not understood. Here, we demonstrate that CtBP2 binds promoters and enhancers in NSCs to directly regulate genes associated with neurogenesis. Surprisingly, our results indicate a prominent role of CtBP2 in activating gene expression in NSCs of the embryonic cortex. This finding is in contrast with the predominantly repressive role of CtBP1/2 in other tissues and cell lines (Stankiewicz et al., 2014). In support of our findings, recent evidence indicates that the *Drosophila* homolog of CtBP1/2 plays important developmental roles as a transcriptional activator (Bi et al., 2022). The mechanisms through which CtBP1/2 activate gene expression in mammalian cells warrant further investigation.

The interaction between PRDM16 and CtBP1/2 plays a key role in brown fat determination by silencing white fat genes (Kajimura et al., 2008). Nonetheless, past reports did not address how the genome-wide occupancy of PRDM16 and CtBP1/2 co-regulates tissue-specific gene expression. Our results suggest that PRDM16 and CtBP2 co-bind enhancers and some promoters in NSCs. In contrast to the predominant role of the PRDM16-CtBP1/2 complex on gene silencing in adipocytes, our results suggest that gene activation through this complex is as prominent as gene repression in the embryonic cortex. In addition, several genes regulated by PRDM16 and CtBP2 in NSCs are associated with neuronal migration. This regulation is expected to be transient, as *Prdm16* is expressed in NSCs and becomes rapidly silenced upon neuronal differentiation (Inoue et al., 2016)(Baizabal et al., 2018). Therefore, our results suggest that transient collaboration between PRDM16 and CtBP1/2 in NSCs influences later stages of neuronal migration.

Our data support a model in which the interaction of PRDM16 with CtBP1/2 mediates the functional cooperation with HDAC1/2, leading to histone deacetylation within regulatory sequences in the developing cortex. Previous work has shown a critical role for HDAC1/2 in the migration and positioning of intermediate cortical progenitors (Tang et al., 2019). Nonetheless, the severe developmental defects observed in *Hdac1/2* double KOs precluded a clear analysis of neuronal migration (Tang et al., 2019). Given the observed genomic overlap between PRDM16, CTBP2, and HDAC1, our data suggests that these factors cooperate in regulating neurogenesis and neuronal migration. Despite their primary role as transcriptional repressors, histone deacetylases also activate gene expression by stimulating transcriptional elongation (Greer et al., 2015). Therefore, HDAC1 might cooperate with PRDM16 and CtBP2 in activating and repressing gene expression in the embryonic cortex. Furthermore, previous reports indicate that CtBP1/2 interact with LSD1 (Ray et al., 2014)(Zeng et al., 2016). Accordingly, we found that LSD1 overlaps with PRDM16 and CtBP2 at a subset of enhancers, suggesting cooperation in gene regulation. This finding is consistent with the role of LSD1 in controlling neuronal migration in the cortex (Zhang et al., 2014). LSD1 promotes migration by activating and repressing gene expression, possibly through H3K9me2 and H3K4me2 demethylation, respectively (Zhang et al., 2014). Hence, our data suggest that PRDM16 collaborates with CTBP1/2, HDAC1/2, and LSD1 in NSCs to control the transcriptional programs of cortical neurogenesis and migration. Further studies are needed to elucidate how the histone methyltransferase activity of PRDM16 regulates cortical enhancers.

In non-neural tissues, PRDM16 and CtBP1/2 are known to be recruited to regulatory sequences via DNA-bound transcription factors (Chi and Cohen, 2016)(Stankiewicz et al., 2014). Our motif enrichment analysis suggests that CTCF is a recruitment factor for CtBP1/2 and their interacting partners. CTCF mediates looping interactions between enhancers and promoters (Dehingia et al., 2022)(Sun et al., 2022). Hence, CtBP1/2 complexes might play a role in promoting or suppressing long-range distal interactions between enhancers and promoters, leading to gene activation or silencing, respectively. The broad expression of the gene encoding CTCF across all tissues suggests that neural-specific transcriptional regulators also participate in recruiting CtBP1/2 to regulatory sequences in the embryonic cortex (Do and Skok, 2024).

*PRDM16* is one of the core genes associated with 1p36 deletion syndrome (Jordan et al., 2015)(Jacquin et al., 2023). Moreover, we recently reported a patient with *PRDM16* haploinsufficiency who exhibited microcephaly, loss of brain gyri, seizures, and intellectual disability (Suresh et al., 2024). The data presented here suggest that PRDM16 interacts with critical regulators of neurogenesis and migration (Tang et al., 2019)(Zhang et al., 2014). Hence, *PRDM16* haploinsufficiency in humans may significantly reduce interactions between PRDM16 and ubiquitous chromatin regulators within transcriptional complexes, potentially contributing to enhancer dysregulation and severe neurodevelopmental deficits.

## LIMITATIONS OF THIS STUDY

Our results suggest that interaction between PRDM16 and CtBP1/2 in NSCs determines subsequent stages of neuronal migration. This model is supported by the observed migration defects after *Prdm16* and *Ctbp1/2* silencing and by the overlap of PRDM16 and CtBP2 binding within sequences that regulate neuronal migration genes. However, CtBP1/2 are also present in cortical neurons. Therefore, we cannot rule out that at least some of the migration defects observed after *Ctbp1/2* silencing *in vivo* are due to gene dysregulation in neurons rather than NSCs. Furthermore, the identification of genes co-regulated by PRDM16 and CtBP2 is still limited by the sensitivity and robustness of Hi-C techniques (Bonev et al., 2017)(Noack et al., 2022)(Ron et al., 2017). For instance, the number of PRDM16-bound enhancers is much larger than the number of EPPs identified by Hi-C in neural progenitors of the embryonic cortex (Baizabal et al., 2018)(Bonev et al., 2017). Hence, we expect that our analysis accounts for only a fraction of the genes directly co-regulated by PRDM16 and CtBP1/2 in NSCs, offering an incomplete picture of the regulatory mechanisms involved in neurogenesis and cell migration.

## MATERIALS AND METHODS

### Mouse strains

All the animal procedures conducted in this study, including housing, husbandry, medical treatment, and euthanasia procedures complied with the protocol approved by the Bloomington Institutional Animal Care and Use Committee (BIACUC) at Indiana University. All mice used in this study for IUE were ICR (CD-1) outbred from Envigo. The conditional *Prdm16^flox/flox^* mice (B6.129-Prdm16^tm1.1Brsp^/J, strain #024992) and the transgenic *Emx1^Cre^*mice (Emx1^tm1(cre)Krj^/J, strain #005628) are available from the Jackson Laboratory and were previously reported (Cohen et al., 2014)(Gorski et al., 2002).

### Knockdown and rescue of *Ctbp1* and *Ctbp2* expression

We used shRNAs that were previously reported to knock down *Ctbp1* and *Ctbp2* efficiently (Siles et al., 2013). The forward sequences are shown below. The shRNA sequences are underlined. *Ctbp1*: 5’-<u>AAACTGTGTCAACAAGGAC</u>TTCAAGAGA<u>GTCCTTGTTGACACAGTTT</u>CTTTTTG-3’ *Ctbp2*: 5’-<u>AAACTGTGTCAACAAAGAA</u>TTCAAGAGA<u>TTCTTTGTTGACACAGTTT</u>CTTTTTG-3’ Scrambled: 5’-<u>GACCTATACGGAGAACAAT</u>TTCAAGAGA<u>ATTGTTCTCCGTATAGGTC</u>CTTTTTG-3’ The shRNA and scrambled sequences were cloned downstream of the U6 promoter of pBluescript plasmid for IUEs. To generate lentiviruses, we cloned the shRNAs and scrambled shRNA into pLL3.7 (Addgene, plasmid #11795). To rescue the knockdown of *Ctbp1/2,* the full-length coding sequences of *Ctbp1* (1326 bp) and *Ctbp2* (2967 bp) were synthesized (GenScript Biotech) and cloned into the EcoRI site of pCIG. The synthesized *Ctbp1* and *Ctbp2* coding sequences carried seven synonymous mutations in the shRNA-targeting sequences. All shRNAs and coding sequences were confirmed by Sanger sequencing.

### *In utero* electroporation

We performed IUE as previously described (Baizabal et al., 2018). Briefly, timed pregnant mice were anesthetized by isoflurane inhalation (3–5%) and received a subcutaneous dose of buprenorphine (0.05–0.1 mg/kg) and Ketoprofen (5 mg/kg) to reduce post-surgery pain. Approximately 1-3 μl of endotoxin-free plasmid dissolved in sterile PBS 1X containing 0.025% Fast Green (SIGMA) was manually injected into the forebrain ventricle of E14.5 embryos. For electroporation, we applied 5 pulses of 30–40 V (50 ms duration and 950 ms intervals) using 7 mm platinum electrodes connected to an ECM 830 square wave electroporator (BTX). A second dose of ketoprofen was administered 24 hrs. post-surgery. To perform the double knockdown of CtBP1 and CtBP2, we co-injected plasmids carrying shRNAs against *Ctbp1* and *Ctbp2* (both at 0.05 μg/μl) together with pCAG-TAG (2.9 μg/μl), which encodes nuclear GFP and membrane TdTomato (Addgene, plasmid #26771). For the rescue experiment, we co-injected *Ctbp1* and *Ctbp2* shRNA plasmids (both at 0.05 μg/μl) with plasmids carrying the shRNA-resistant *Ctbp1* and *Ctbp2* coding sequences (both at 0.1 μg/μl), and pCAG-TAG (2.7 μg/μl).

### Primary cultures of cortical NSCs

E14.5 cortices from WT and *Prdm16* KO embryos were dissected and incubated for 10 min at 37°C in 3 ml of 0.05% trypsin with EDTA (Gibco, 25300062) and 0.01% DNase (Sigma, DN-25). Trypsin was removed and the cortices were washed twice with 8 ml of Hibernate-E minus calcium (BrainBits, HECA500). After the last wash, 3 ml of dissociation solution consisting of Hibernate-E minus calcium plus 0.01% DNase was added to the cortices and dissociated into a single-cell suspension using a fired-polished Pasteur pipette. Cells were spun at 1000 rpm for 5 min at room temperature (RT) and resuspended in complete NSC media consisting of Dulbecco’s Modified Eagle Medium/Nutrient Mixture F12 (DMEM/F12; Gibco, 11320033), 1X GlutaMax supplement (Gibco, 35050-061), 1X B27 supplement, 10 ng/ml murine Epidermal Growth Factor (EGF; PeproTech, 315-09), 10 ng/ml human Fibroblast Growth Factor 2 (FGF2; PeproTech, 100-18), and 0.12 mg/ml bovine serum albumin (Gibco, 15260037). Cells were plated on plates pre-treated with 10 μg/ml Poly-D-Lysine (Corning, 354210) and 12 μg/ml Laminin (Gibco, 23017015). Half the media was replaced every two days. For subcultures, cells were treated with 0.05% trypsin plus EDTA, washed once with NSC media and plated on freshly prepared Poly-D-Lysine and Laminin pre-treated plates.

### Lentivirus and retrovirus generation and infection

Confluent HEK 293T cells were transfected with lentiviral vectors using lipofectamine. To transfect cells in a 10 cm petri dish, the following components were added to 3 ml Opti-MEM I with GlutaMAX (Gibco, 51985034): 10 μg pLL3.7 (Addgene, plasmid #11795), 5 μg pRRE (Addgene, plasmid #12251), 2.5 μg pRSV-Rev (Addgene, plasmid #12253), 2.5 μg pMD2.G (Addgene, plasmid #12259), 41 μl Lipofectamine 3000 transfection reagent, and 35 μl P3000 Enhancer Reagent (Invitrogen, L3000015). The supernatant of transfected cells was collected after 24 and 48 hrs., pooled into a single 50 ml Falcon tube, and centrifuged at 2000 rpm for 10 min at RT to remove cellular debris. The supernatant was filtered using a 45 μm pore-size filter and laid on a 10% sucrose buffer (4 volumes supernatant per 1 volume sucrose buffer). The sucrose buffer consisted of 10 g/dL sucrose, 100 mM NaCl, 50 mM Tris-HCl, and 0.5 mM EDTA. All components of the sucrose buffer were diluted in water, adjusted at pH 7.4, and filter-sterilized. The viral supernatant laying on top of 10% sucrose buffer was then centrifuged at 10,000 g for 4 hrs. at 4°C to concentrate the lentiviral particles. The supernatant was discarded without disturbing the pellet. The pellet was resuspended in 100 μl of PBS 1X and stored in aliquots at −80°C. Confluent NSCs were co-infected by directly adding lentiviruses carrying shRNAs for *Ctbp1* and *Ctbp2*, each to a final 1:100-1:200 dilution. To increase infection efficiency, we preferentially used freshly prepared lentiviruses.

Retroviruses carrying the full-length coding sequence of *Prdm16* or mutant *Prdm16* were also produced by co-transfecting HEK 293T cells with MSCV PRDM16 expression vector (Addgene, plasmid #15504), pBS-CMV-gag pol (Addgene, plasmid #35614), and pMD2.G. The MSCV PRDM16 vector was used as a template to produce the mutant *Prdm16* by changing the GACCTG sequence to GCAAGC using the QuikChange II XL Site-Directed Mutagenesis Kit (Agilent Technologies #200521) following the manufacturer’s instructions. This mutation changes the amino acid sequence of the PLDLS motif to PLASS, which completely blocks the interaction between PRDM16 and CtBP1/2 (Kajimura et al., 2008).

### Immunoprecipitation and western blot

To increase the sensitivity of our IP assays, we infected cortical NSCs with a retrovirus expressing the full-length coding sequence of *Prdm16*. In addition, we immunoprecipitated the endogenous CtBP1/2 to confirm their interaction with PRDM16.

#### Nuclear extraction

Flash-frozen cell pellets were thawed on ice and resuspended in 1 ml of cold hypotonic Buffer A (10 mM HEPES pH 7.5, 1 mM MgCl₂, 10 mM KCl, 0.5 mM DTT) supplemented with 1X protease inhibitor (Thermo Scientific, 87786). The suspension was transferred to a pre-cooled Eppendorf tube and gently ground on ice with a sterile pestle. The sample was then centrifuged at 1500 g for 5 minutes at 4°C. The resulting nuclei pellet was resuspended in 1 ml of pre-cooled nuclear extraction Buffer B (20% glycerol, 20 mM HEPES, pH 7.3, 250 mM NaCl, 1.5 mM MgCl₂, 0.2 mM EDTA, 0.5 mM DTT, and 1X protease inhibitor). The nuclei suspension was then homogenized on ice using a sterile pestle until a cloudy appearance was observed, indicating chromatin release from broken nuclei. The homogenate was incubated for 30 minutes at 4°C with mixing and centrifuged at maximum speed for 10 minutes at 4°C. The supernatant, containing the nuclear extract, was collected.

#### Bead preparation

Protein G Dynabeads (Invitrogen, 10003D) were washed twice in 1 ml PBS 1X plus 0.1% Tween 20. Per IP reaction, 6 µg of antibodies were added to 50 µl of washed protein G Dynabeads and incubated for 30 minutes at RT with rotation. The antibody solution was discarded and the beads were washed twice with conjugation buffer (100 mM sodium tetraborate, pH 9.0). To crosslink Protein G dynabeads to antibodies, 500 µl of 20 mM dimethyl pimelimidate (DMP, Thermo Scientific, 21666) diluted in conjugation buffer were combined with 50 µl of antibody-conjugated beads and incubated in the dark at RT for 30 minutes with rotation. The DMP solution was discarded and the beads were washed twice with 1 ml quenching buffer (200 mM Tris, pH 8.0). After the second wash, 1 ml of quenching buffer was added and the beads were incubated at RT for 45 minutes while mixing. Finally, the quenching buffer was aspirated and the beads were washed once with Buffer B before being resuspended in 100 µl of the same buffer.

#### Immunoprecipitation

100 µl of antibody-conjugated beads in buffer B were added to 1 ml of nuclear extract. The mixture was incubated overnight at 4°C with rotation. After incubation, the beads were washed once with 1 ml of Buffer B for 10 minutes at 4°C with rotation. The beads were then washed twice with 1 ml of high salt wash buffer (20 mM HEPES pH 7.3, 500 mM NaCl, 1.5 mM MgCl_2_, 0.2 mM EDTA, 10% glycerol, 0.1% NP40) for 10 minutes at 4°C with rotation, followed by two washes with 1 ml of low salt wash buffer (20 mM HEPES pH 7.3, 150 mM NaCl, 1.5 mM MgCl_2_, 0.2 mM EDTA, 10% glycerol, 0.1% NP40) under the same conditions. To elute immunoprecipitated proteins, the beads were incubated with 50 µl elution buffer (0.1 M glycine, 0.1% NP40 pH 2.5, and 1X protease inhibitor) for 40 minutes at RT with rotation. Finally, 1/10 volume of 1 M Tris was added to neutralize the sample, which was then used for WB. To prepare samples for proteomics, the beads were washed twice with 1 ml of low salt wash buffer as indicated above, followed by two additional washes with 1 ml of low salt wash buffer without glycerol and NP40. The proteins were then digested on the beads and identified by mass spectrometry.

#### Western blot

The eluted samples were boiled with Laemmli sample buffer (Bio-Rad) and loaded into 4–20% mini-Protean TGX precast protein gels (Bio-Rad). Protein was transferred to PVDF membranes and incubated in blocking buffer (3% Bovine Serum Albumin, 20 mM Tris-HCl pH 7.5, 150 mM NaCl, and 0.1% Tween 20) for 1 hr. at RT. The membranes were then incubated with primary antibodies overnight at 4°C, washed 3 times, incubated with secondary antibodies for 1 hr. at RT, and washed 3 times. The following antibodies were used: sheep anti-PRDM16 (R&D Systems, AF6295, 1:1000), rabbit anti-PRDM16 (gift from Patrick Seale, UPenn), sheep IgG control antibody (Millipore, 12-370), anti-CtBP1 (Proteintech, 10972-1-AP, 1:1000), anti-CtBP2 (Invitrogen, PA5-86439, 1:1000), anti-HDAC1 (Diagenode, C15410325, 1:1000), anti-HDAC2 (Diagenode, C15200201, 1:1000), secondary goat anti-rabbit IgG (LICORbio, 926-32211, 1:5000 dilution), secondary anti-sheep IgG HRP-conjugated (R&D Systems, HAF016, 1:1000). For HRP-conjugated antibodies, we used Clarity Western ECL Substrate (BioRad, 1705060]. Gels were imaged with an Odyssey CLx Imager (LICORbio) or ChemiDoc MP imaging system (BioRad).

### Immunostaining

P5 mice were anesthetized with Xylazine (20 mg/Kg) - Ketamine (150 mg/Kg) and their brains were perfused with 4% paraformaldehyde (PFA) diluted in PBS 1X. Perfused brains were washed with PBS 1X three times and sliced into 75–100 μm sections using a VT1000 Vibratome (Leica Biosystems). Antigen retrieval was performed by incubating the tissue sections in a solution containing 10 mM sodium citrate and 0.05% Tween 20 pH 6.0 for 1 hr. at 70°C. Tissue sections were rinsed in PBS 1X three times and incubated for 1 hr. in blocking buffer (10% goat serum, 0.1% Triton X-100, 0.01% sodium azide diluted in PBS 1X) at RT with mixing. Primary antibodies diluted in the blocking buffer were added to the tissue sections and incubated overnight at 4°C with mixing. Brain sections were washed three times with PBS 1X and incubated for 2 hrs. at RT with Alexa Fluor-conjugated secondary antibodies diluted in blocking buffer. Sections were washed again three times with PBS 1X and nuclei were stained with DAPI (4’,6-diamidino-2-phenylindole) before mounting with Fluoromount-G (Southern Biotech). We used the following primary antibodies and dilutions: anti-CtBP1 (Proteintech, 10972-1-AP, 1:100), anti-CtBP2 (Invitrogen, PA5-86439, 1:100), anti-BRN2 (Santa Cruz Biotechnology, sc-393324, 1:200), anti-GFP (Aves Lab, GFP-1020, 1:4000), anti-Ki67 (Abcam, ab16667, 1:250), and anti-Nestin (R&D Systems, MAB2736, 1:500). To detect primary antibodies, we used anti-rabbit and anti-mouse secondary antibodies conjugated to Alexa Fluor 488, 546, or 647 (Invitrogen, 1:500). Imaging was performed with a Leica SP8 confocal microscope and processed with the LAS X Life Science microscope software (Leica Microsystems).

### RNA sequencing

Total RNA was extracted using the RNeasy kit (Qiagen) following the manufacturer’s instructions. The samples consisted of four independent cultures of NSCs infected with lentiviruses carrying either scrambled shRNA or *Ctbp1/2* shRNAs. The quality of the RNA samples was assessed using TapeStation 2200 (Agilent). We performed quantitative PCR in all replicates to confirm efficient *Ctbp1/2* knockdown ahead of sequencing. The RNA-seq libraries were prepared by Illumina Stranded mRNA Library Prep protocol (Illumina, 20040534) and analyzed with TapeStation 4200 (Agilent). The libraries were pooled and loaded on a NextSeq 1000/2000 P2 (100 Cycles) v3 flow cell (Illumina, 20046811) configured to generate 2×59 nucleotides paired-end reads. The demultiplexing of the reads was performed using bcl2fastq2 Conversion Software version 2.20.

### Chromatin immunoprecipitation and sequencing

E14.5 cortices from an entire litter were dissociated and pooled into a single-cell suspension consisting of approximately 20-40 million cells, which is sufficient for one transcription factor ChIP-seq. Samples were dual crosslinked by incubating in 1.5 mM EGS (ethylene glycol bis[succinimidyl succinate]) solution (Thermo Scientific) for 20 min followed by 1% PFA for 10 min at RT. Crosslinking was quenched with 125 mM glycine and cells were washed twice with cold PBS 1X plus protease inhibitor (Thermo Scientific, 87786). Cells were centrifuged and resuspended in lysis buffer (20 mM Tris-HCl pH 8.0, 85 mM KCl, 0.5% NP40). Samples were centrifuged at 1500 g for 5 min and the nuclei were resuspended in SDS buffer (0.2% SDS, 20 mM Tris-HCl pH 8.0, 1 mM EDTA). Nuclei were sonicated with an S220 ultrasonicator (Covaris) to generate 100-500 bp chromatin fragments. After spinning chromatin at 18,000 g for 10 min, the supernatant was transferred to a clean tube and one volume of 2X ChIP dilution buffer (0.1% sodium deoxycholate, 2% Triton X-100, 2 mM EDTA, 30 mM Tris-HCl pH 8.0, 300 mM NaCl) was added. Around 1% of the sample was set aside as input and the remaining supernatant was incubated overnight at 4°C with 5 µg of anti-CtBP2 antibody (Invitrogen, PA5-86439), anti-HDAC1 antibody (Diagenode, C15410325), or anti-LSD1 antibody (Abcam, ab17721). The next day, 50 µL of Protein G Dynabeads (Invitrogen, 10003D) were added and incubated for 2 hrs. at 4°C. After incubation, beads were washed twice with low salt wash buffer (0.1% SDS, 1%Triton X-100, 2 mM EDTA, 20 mM Tris-HCl pH 8.0, 150 mM NaCl), twice with high salt wash buffer (0.1% SDS, 1% Triton X-100, 2 mM EDTA, 20 mM Tris-HCl pH 8.0, 500 mM NaCl), twice with LiCl wash buffer (0.25 M LiCl, 0.5% NP40, 0.5% sodium deoxycholate, 1 mM EDTA, 10 mM Tris-HCl pH 8.0), and twice with TE pH 8.0 (10 mM Tris-HCl, 1 mM EDTA). Chromatin was eluted from beads using elution buffer (1% SDS, 0.1 M NaHCO3) and the recovered supernatant was incubated in reverse crosslinking solution (250 mM Tris-HCl pH 6.5, 62.5 mM EDTA pH 8.0, 1.25 M NaCl, 5 mg/ml of Proteinase K) at 65°C overnight. DNA was then extracted with phenol/chloroform/isoamyl alcohol, precipitated with 3M sodium acetate pH 5.0, and resuspended in TE pH 8.0 low EDTA (10 mM Tris-HCl, 0.1 mM EDTA). Finally, samples were treated with RNase A (100 mg/ml) for 30 min at 37°C. For quantitative CtBP2 ChIP-seq, 120 ng of spike-in chromatin (Active Motif, 53083) and 2 µg of spike-in antibody (Active Motif, 61686) were included in the protocol. For library preparation, we used the NEBNext Ultra II DNA Library Prep kit (NEB, E7103S) with NEBNext Multiplex Oligos for Illumina (NEB, E7335S). The ChIP-seq libraries were loaded on a P2 (100 Cycles) flow cell configured to produce 2×61 nucleotides paired-end reads and sequenced on a NextSeq 2000 instrument.

### Sequencing analyses

#### RNA-seq analysis

Raw reads were subjected to quality control using FastQC. The raw reads were trimmed using Trimmomatic (Bolger et al., 2014). rRNA contaminations were removed using TagDust2 (Lassmann, 2015). The cleaned raw reads were then examined again using FastQC. The trimmed raw reads were aligned to the mm39 (BSgenome.Mmusculus.UCSC.mm39chrs.fa) reference genome using the STAR aligner with the “--alignEndsType EndToEnd” parameter since the raw reads were trimmed (Dobin et al., 2013). Gene counting was performed in the same STAR alignment script by adding “--quantMode GeneCounts”. Differential gene expression analysis on the raw counts matrix was performed using DESeq2 (Love et al., 2014). The volcano plot was generated using ggplot2 from the Tidyverse package. The heat map was generated using the ComplexHeatmap package (Gu et al., 2016). The RNA-seq raw reads from *Prdm16* WT and KO cortices were downloaded using the fasterq-dump command from the SRA tools kit software (Sayers et al., 2022). Gene ontology analyses were performed using PANTHER19.0 (Mi et al., 2019).

#### ChIP-seq analysis

The quality of ChIP-seq raw reads was checked using FastQC. The ChIP-Seq raw reads were trimmed using Trimmomatic and aligned to the mm39 (BSgenome.Mmusculus.UCSC.mm39chrs.fa) reference genome using the Bowtie2 aligner with the “--no-unal” parameter (Langmead and Salzberg, 2012). The outputted SAM files were converted to BAM files using SAMtools (Danecek et al., 2021). The bigWig files were generated using the bamCoverage command from deepTools using “--binSize 10 --smoothLength 30 – centerReads” parameters (Ramírez et al., 2016). The peaks were called using MACS3 with a q-value threshold of 0.05, using “--narrow” for the transcription factor ChIP-Seq samples and “--broad” for the histone ChIP-Seq samples (Zhang et al., 2008). The “exclude ranges” (ChIP-seq blacklisted regions) were compiled in “mm39.excluderanges.bed” and removed from the BED files using the BEDTools intersect command with the “-v” argument (Ogata et al., 2023)(Quinlan and Hall, 2010). The ChIP-Seq raw reads for H3K27ac, H3K4me1, H3K4me3, CTCF, and ATAC-seq raw reads from previously published papers were downloaded using the fasterq-dump command from the SRA tools kit software (Baizabal et al., 2018)(Stamatoyannopoulos et al., 2012)(Yue et al., 2014)(Sayers et al., 2022).

The reproducibility of ChIP-Seq peaks was tested by intersecting the peaks of replicates 1 and 2 via the BEDTools intersect command with the “-f 0.3 -r” parameters. The intersection of different ChIP-seq datasets was done via the intersect command with the “-f 0.5 -F 0.5 -e” parameters. The spike-in CtBP2 ChIP-Seq was mapped to a composite *Mus musculus*-*Drosophila melanogaster* reference genome (mm39-dm6.fa) via Bowtie2. The scaling factor was based on the Active Motif protocol (https://www.activemotif.com/documents/1977.pdf). The reads mapped to mm39 were moved to a separate BAM file and the bigWigs were generated with the calculated scaling factor using deepTools bamCoverage (Ramírez et al., 2016). The ChIP-Seq enrichment matrices were generated using deepTools computeMatrix for the heatmaps and profile projections (Ramírez et al., 2016). The ChIP-Seq heatmaps were generated using deepTools plotHeatmap and the profile projections were generated using deepTools plotProfile (Ramírez et al., 2016). HOMER was utilized to annotate peaks and perform motif analysis (Heinz et al., 2010).

#### Identification of distally regulated genes

Previously published Hi-C datasets from the mouse embryonic cortex were used to identify genes directly regulated by a subset of enhancers in neural progenitors (Bonev et al., 2017). The enhancer coordinates of EPPs and their associated genes are listed in their supplementary table under the “NPC_based” tab (https://ars.els-cdn.com/content/image/1-s2.0-S0092867417311376-mmc3.xlsx). The reported enhancer coordinates represent the center of 5 Kb intervals. In our analysis, each enhancer coordinate was extended by 2.5 Kb in both directions. The resulting coordinates were converted from mm10 to mm39 using LiftOver (https://genome.ucsc.edu/cgi-bin/hgLiftOver) (Hinrichs et al., 2006). We confirmed a high enrichment of H3K27ac in the resulting Hi-C mm39 coordinates, indicating that the majority of these genomic regions are enhancers in the embryonic cortex.

### Reproducibility

To ensure the complete reproducibility of our bioinformatics analyses, all third-party software packages used in this project were installed in an apptainer/singularity container (Kurtzer et al., 2017) available for download at https://BrendelGroup.org/SingularityHub/PRDM16_CtBP_Project.sif. Workflow and analysis scripts and complete documentation of the computational work are accessible on GitHub: https://github.com/BrendelGroup/PRDM16_CtBP_Project. The workflow scripts will download all data analyzed and re-generate all the analysis results described in the paper. The modular design of the workflows also allows readers to repurpose the scripts to analyze their data similarly.

### Statistical analyses

#### Western blot quantification

Western blot bands were quantified using ImageJ (National Institutes of Health, USA). Gel images were converted to 16-bit files and the rectangular selection tool was used to define the band areas. Profile plots were generated using the Gel analyzer tool to calculate the area under each peak, representing the relative band intensity. The relative band intensities of CtBP1 and CtBP2 were normalized against α-Tubulin to determine knockdown efficiency.

#### Quantification of cell migration

Electroporated cortices were collected from one litter for the control condition and two litters for the knockdown and rescue conditions. To quantify cell migration, a minimum of six immunostained sections from three brains (control condition) or four brains (knockdown and rescue conditions) were used to manually count 80-300 GFP^+^ cells using ImageJ/Fiji. Imaging settings were identical between the experimental and control groups. The proportion of GFP^+^ cells in deep layers over the total number of GFP^+^ cells was quantified for each condition. The statistical significance was determined using the two-tailed unpaired Student’s t-test with the Prism Software (GraphPad) and reported as *p<0.05, **p<0.01, ***p<0.001, ****p<0.0001.

## ACKNOWLEDGEMENTS

We thank Justin Kumar and Heather Hundley (both at Indiana University Bloomington) for their valuable comments and recommendations during the development of this project. We thank the Center for Genomics and Bioinformatics for their support in next-generation sequencing experiments. This work was supported by Indiana University startup funding to J-M.B.

## AUTHOR CONTRIBUTIONS

**Sophie Warren:** Methodology, Validation, Formal Analysis, Investigation, Writing - Review & Editing, Visualization. **Bader El Farran:** Software, Validation, Formal Analysis, Investigation, Writing - Review & Editing, Visualization. **Sungyun Kang:** Methodology, Validation, Investigation. **Adhyeta Choudhuri:** Validation, Investigation, Writing - Review & Editing. **Sen Xiong:** Investigation. **Volker P. Brendel:** Software, Formal Analysis, Data Curation, Writing - Review & Editing, Supervision. **José-Manuel Baizabal:** Conceptualization, Investigation, Writing - Original Draft, Supervision, Project Administration, Funding Acquisition.

## DATA AVAILABILITY

The raw sequencing data generated in this study have been deposited in the NCBI Sequence Read Archive (SRA) under accession number PRJNA1134662.

---

**Fig. S1.**
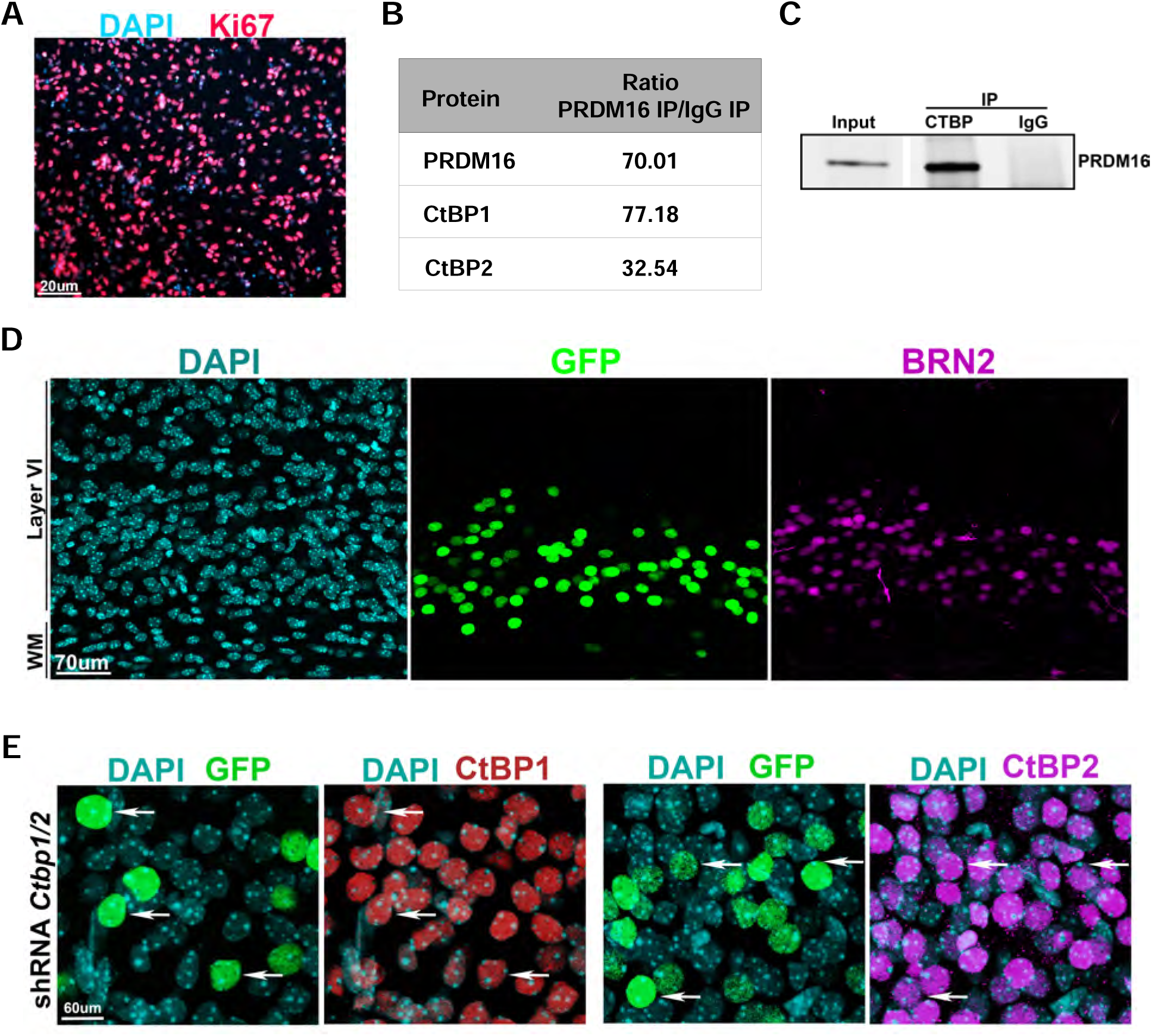
(A) Cortical NSCs in culture are positive for the proliferation marker Ki67. (B) Immunoprecipitation (IP) of PRDM16 followed by mass spectrometry analysis indicates a strong interaction with CtBP1 and CtBP2 in cortical NSCs *in vitro*. (C) IP of CtBP1/2 confirms interaction with PRDM16 in NSCs *in vitro*. An unspecific IgG was used as a negative control. (D) *In-utero* electroporation of *Ctbp1/2* shRNAs and *GFP* at E14.5 results in ectopic GFP^+^/BRN2^+^ upper-layer neurons in layer VI of the P5 cortex. (E) Co-electroporation of *Ctbp1/2* shRNAs and *GFP* with plasmids encoding the shRNA-resistant coding sequences of *Ctbp1* and *Ctbp2* results in the rescue of CtBP1 and CtBP2 levels in cortical neurons (arrows).

**Fig. S2.**
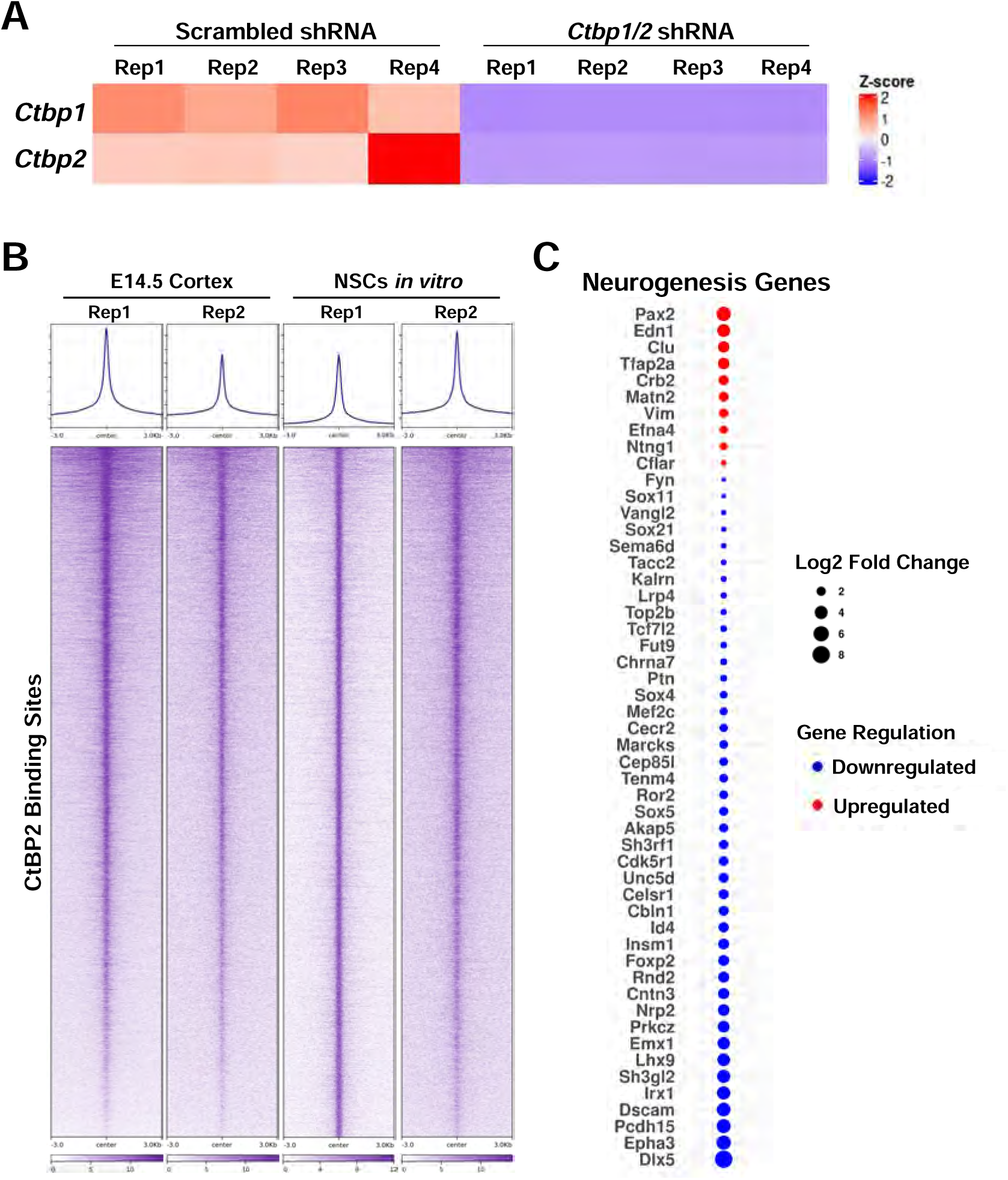
(A) Heatmap showing *CtBP1/2* relative expression in four RNA-seq biological replicates (Rep) per condition. The heatmap confirms *Ctbp1/2* silencing in NSCs infected with lentiviruses carrying shRNAs in comparison to lentiviruses carrying scrambled shRNA controls. (B) Heatmaps showing CtBP2 binding in the E14.5 cortex and NSCs *in vitro*. Two replicates are shown for each sample type. (C) The dot plot indicates neurogenesis genes directly regulated by CtBP2 that display differential expression in the *Ctbp1/2* knocked down NSCs.

**Fig. S3.**
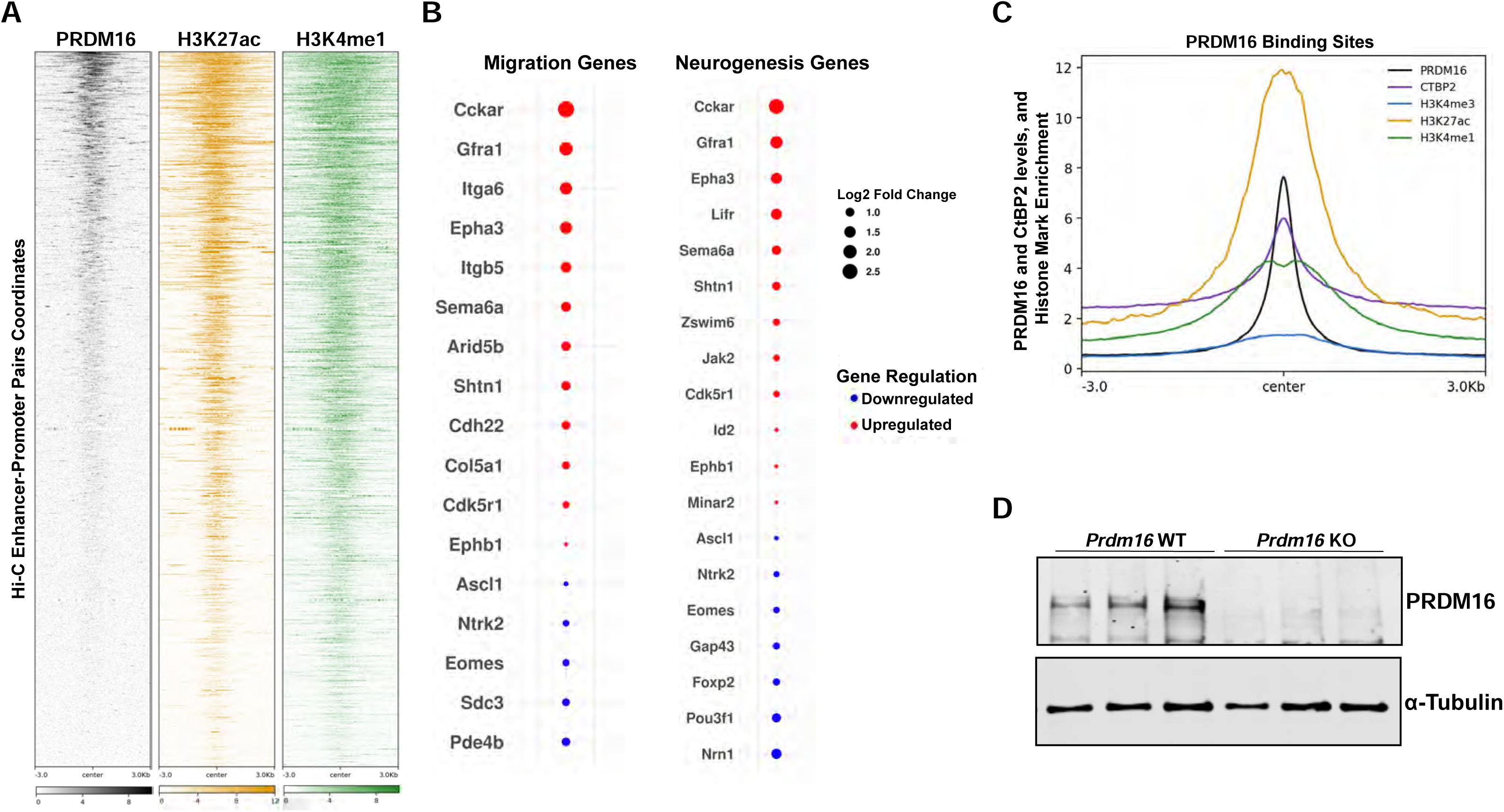
PRDM16, H3K27ac, and H3K4me1 levels within the enhancer regions of all enhancer-promoter pairs identified by Hi-C in neural progenitors of the embryonic cortex (Bonev et al., 2017). (B) Dot plot of migration and neurogenesis genes predicted to be directly regulated by PRDM16 that display differential expression in NSCs (PAX6^+^ cells) of the *Prdm16* E15.5 KO cortex (Baizabal et al., 2018). (C) The projection profiles indicate the levels of PRDM16, CtBP2, and histone marks within PRDM16-regulated sequences. (D) *Prdm16* WT and KO cultures of NSCs were generated from E14.5 cortices. The western blot confirms PRDM16 depletion in the KO cells. Each lane in the gel represents an independent NSC line derived from an individual cortex. The loading control is α-Tubulin.

**Fig. S4.**
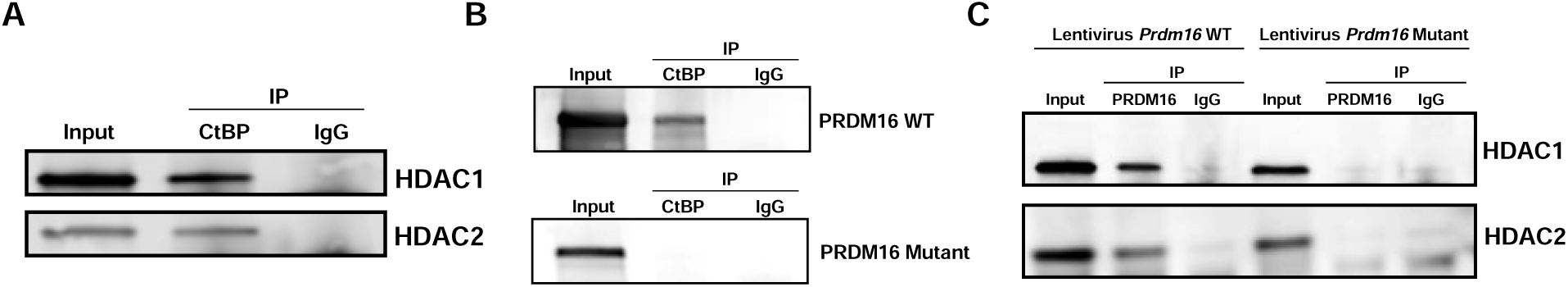
(A) Co-immunoprecipitation (IP) of CtBP1/2 (collectively referred to as CtBP in this figure) reveals an interaction with HDAC1 and HDAC2 in cortical NSCs *in vitro*. (B) 293T cells were transfected with plasmids encoding either WT *Prdm16* or mutant *Prdm16* that is predicted not to bind CtBP1/2. Two days after transfection, CtBP1/2 were immunoprecipitated, and western blot analysis confirmed an interaction with the WT but not the mutant PRDM16. (C) *Prdm16* KO NSCs *in vitro* were infected with lentiviruses expressing either WT *Prdm16* or mutant *Prdm16*. The WT PRDM16 interacts with HDAC1 and HDAC2, whereas the mutant PRDM16 does not, suggesting that these interactions depend on PRDM16 binding to CtBP1/2. An unspecific IgG was used as a negative control in all panels.

